# Optogenetic control of cell signaling with red/far-red light-responsive optogenetic tools in *Caenorhabditis elegans*

**DOI:** 10.1101/2022.08.12.503710

**Authors:** Shigekazu Oda, Emi Sato-Ebine, Akinobu Nakamura, Koutarou D. Kimura, Kazuhiro Aoki

**Author notes:** Corresponding contributor. These authors equally contributed to this work (Shigekazu Oda, Emi Sato-Ebine).

## Abstract

Optogenetic techniques have been intensively applied to the nematode *Caenorhabditis elegans* to investigate its neural functions. However, as most of these optogenetics are responsive to blue light and the animals exhibits avoidance behavior to blue light, the application of optogenetic tools responsive to longer wavelength light has been eagerly anticipated. In this study, we report the implementation in *C. elegans* of a phytochrome-based optogenetic tool that responds to red/near-infrared light and manipulates cell signaling. We first introduced the SynPCB system, which enabled us to synthesize phycocyanobilin (PCB), a chromophore for phytochrome, and confirmed the biosynthesis of PCB in neurons, muscles, and intestinal cells. We further confirmed that the amount of PCBs synthesized by the SynPCB system was sufficient for photoswitching of phytochrome B (PhyB)-phytochrome interacting factor 3 (PIF3). In addition, optogenetic elevation of intracellular Ca^2+^ levels in intestinal cells induced a defecation motor program. These SynPCB system and phytochrome-based optogenetic techniques would be of great value in elucidating the molecular mechanisms underlying *C. elegans* behaviors.

## Introduction

*Caenorhabditis elegans*, a multicellular organism, is widely used in a wide range of research fields, because of its ease of culture, short life cycle, and the availability of genetic information ^1^. Genetic approaches using *C. elegans* have yielded fruitful results in the discovery of genes related to various cellular and physiological functions. In general, however, there are limitations to the genetic approach when lethal or redundant genes are targeted, and phenotypes produced by genetic alteration are masked by adaptation through a feedback mechanism. To address these issues, it is necessary to apply acute perturbations and observe the changes at a sampling rate faster than the characteristic time constant of the dynamic system ^2,3^. Although a drug-inducible expression system is a possible tool, it takes a longer time than the typical behaviors observed in nematodes. An alternative method using small compounds is not suitable for perturbation at a specific tissue. Therefore, there is a need for tools to manipulate cellular functions in *C. elegans* with higher temporal and spatial resolution.

Optogenetics is a powerful technique for controlling the function of proteins, cells, and individuals by ectopically expressing light-responsive proteins ^4^. Light-gated ion channels, particularly channelrhodopsin, have long been applied to *C. elegans* since the very early days of the method ^*5,6*^. Light-induced dimerization (LID) systems, which alter protein-protein interactions in response to light, enable the manipulation of intracellular cell signaling ^7,8^. Among them, blue light-responsive LID systems such as CRY2/CIB and iLID/SspB have also been employed in *C. elegans* for a variety of applications ^9–12^. *C. elegans* exhibits avoidance behavior (negative phototaxis) to short wavelengths of light, especially UV light, through LITE-1 in a group of photosensory neurons ^13^. To circumvent this issue, a *C. elegans* mutant lacking *LITE-1* is often used when using the blue light-responsive optogenetic tools. However, the extent to which *LITE-1* mutation affects the physiological functions of *C. elegans* must be carefully considered. Therefore, the introduction of optogenetic tools that do not use blue light to *C. elegans* has been anticipated.

The phytochrome B (PhyB)/phytochrome interacting factor (PIF) is one of the LID tools that respond to red light/far-red light ^14,15^; red light induces the binding of PhyB to PIF, and far-red light causes the dissociation of the PhyB-PIF complex (Fig. 1A). PhyB requires chromophores such as phytochromobilin (PΦB) and phycocyanobilin (PCB) for the photoreception. These chromophores are covalently attached to PhyB. Because PΦB and PCB are found only in plants and cyanobacteria, it is necessary to purify and add chromophores to cells ectopically expressing the PhyB/PIF system. Several groups, including ours, have succeeded in developing genetically encoded systems for the synthesis of PCB in mammalian cultured cells ^16–18^. Our PCB synthesis system (hereinafter SynPCB system) is composed of four genes required for PCB synthesis, namely, the genes encoding PCB:ferredoxin oxidoreductase (PcyA), heme oxygenase1 (HO1), ferredoxin (Fd), and Fd-NADP+ reductase (Fnr) (Fig. 1A). These genes are expressed in the mitochondrial matrix where heme is abundant. We have recently reported that the SynPCB2.0 and SynPCB2.1 systems, which are improved versions of the SynPCB system ^19^, allow efficient synthesis of PCB in mammalian cells, fission yeast, and *Xenopus* embryo. However, there are no reports on whether the PhyB-PIF LID system is applicable to *C. elegans*.

**Figure 1.**
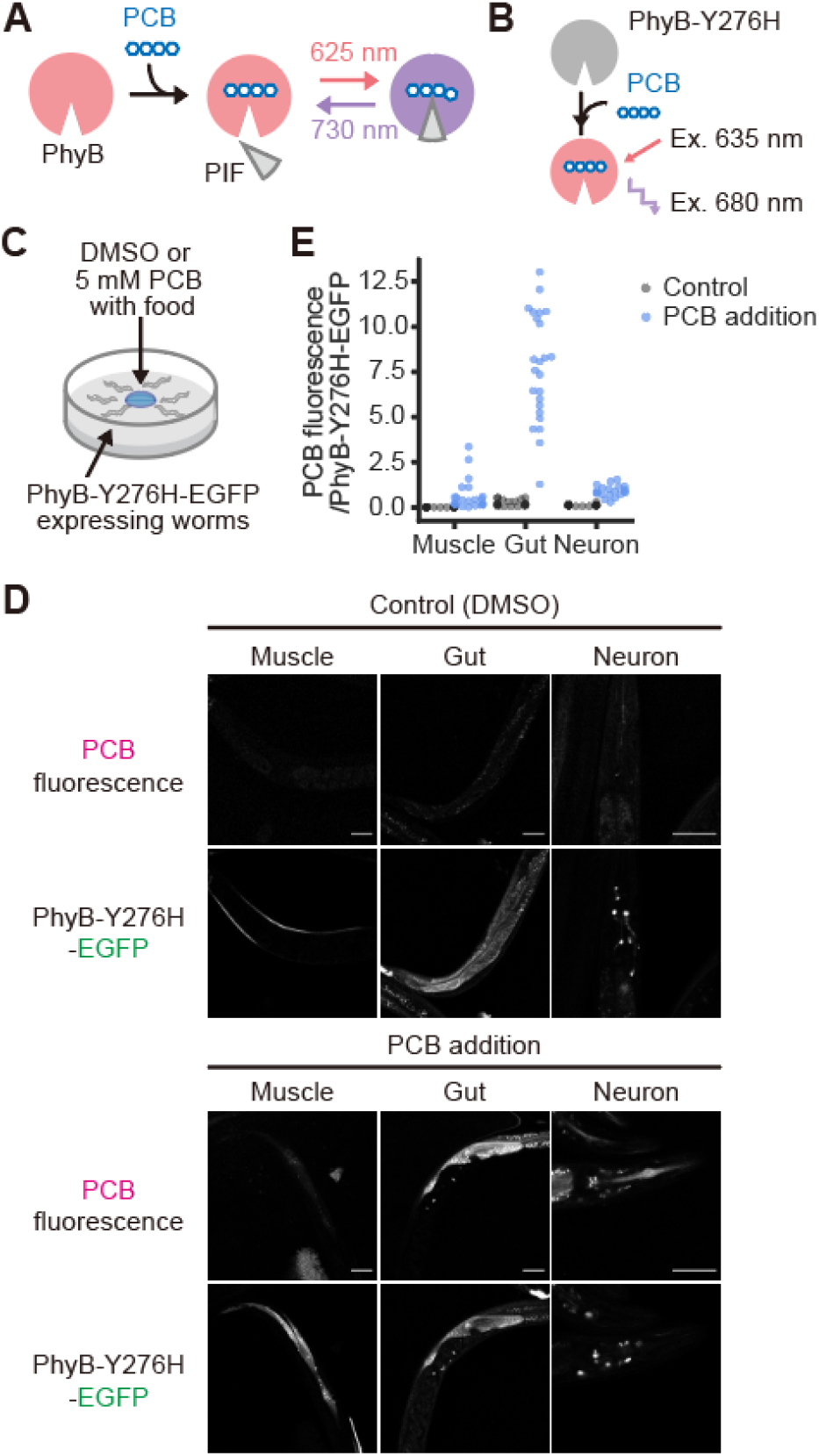
Delivery of purified PCB to the tissues in *C. elegans*. (A) Schematic representation of the PhyB-PIF light switch. Apo-PhyB protein (left) incorporates PCB, a chromophore, to generate holo-PhyB. Upon red-light illumination, the holo-PhyB binds to PIF. Far-red light exposure induces dissociation of the PhyB-PIF complex. (B) PhyB-Y276H mutants emit near-infrared fluorescence when PhyB-Y276H binds to PCB. (C) To examine whether PCB is delivered to the tissues in *C. elegans*, worms expressing PhyB-Y276H-EGFP in the muscle, gut, or neurons were cultured with or without 5 mM PCB overnight. (D) Representative images of PCB (upper row) and PhyB-Y276H-EGFP (lower row) are shown in control (upper panels) and PCB-treated worms (lower panels). Scale bars, 60 μm. (E) The ratio values of PCB fluorescence to PhyB-Y276H-EGFP fluorescence are plotted under the indicated conditions. Control: N = 18 (muscles), 20 (gut), and 19 (neurons). PCB addition: N = 21 (muscles), 24 (gut), 20 (neurons).

In this study, we demonstrated that the SynPCB system allows synthesis of a sufficient amount of PCB for efficient PhyB/PIF photoswitching by light in *C. elegans*. We found that the purified PCB could only be delivered to the intestine of *C. elegans* and could not reach deeper tissues such as the neurons and muscles. Meanwhile, the introduction of SynPCB in *C. elegans* yielded PCB synthesis in the intestinal, neurons, and muscles. Next, we confirmed the light-induced change in the subcellular localization of PIF proteins through binding to PhyB. Finally, we manipulated the intracellular cell signaling and behavior of *C. elegans* with the PhyB-PIF system.

## Results

### Limited intracellular delivery of purified PCB in *C. elegans*

We first investigated whether purified PCB could also be used in *C. elegans*, because the addition of PCB purified from cyanobacteria has enabled the use of the PhyB/PIF LID system in budding yeast, cultured cells, and zebrafish ^15,20–22^. For this purpose, we used a PhyB mutant (PhyB-Y276H), which emits near-infrared fluorescence when bound to PCB (Fig. 1B). Worms expressing PhyB-Y276H-EGFP in the intestines, neurons, or muscles were established and fed with DMSO or 5 mM PCB solution overnight (Fig. 1C). No near-infrared fluorescence was observed in any tissues of the DMSO-fed animals (Fig. 1D and 1E). On the other hand, in animals fed PCB, near-infrared fluorescence was observed in the intestine, but not in the neurons or muscles (Fig. 1D and 1E). These results indicate that purified PCB can reach the intestine but not the deeper tissues such as nerves and muscles in *C. elegans*.

### The SynPCB system synthesizes PCB in the intestine, neurons, and muscles of *C. elegans*

Next, we applied the SynPCB, a genetically encoded PCB synthesis system, to *C. elegans*. Cyanobacteria produce PCB by metabolizing heme and biliverdin through four responsible proteins, PcyA, HO1, Fd, and Fnr (Fig. 2A). Previous studies have shown that PCBs can beefficiently synthesized in cultured cells by expressing these four genes in the mitochondria ^17,18^. Recently, we succeeded in improving the genetically encoded PCB synthesis systems, SynPCB2.0 and SynPCB2.1, which are polycistronic gene expression systems in which PcyA, HO1, and Fnr-Fd chimeric proteins with a mitochondrial transition signal (MTS) are linked by the self-cleaved P2A peptides ^19^.

**Figure 2.**
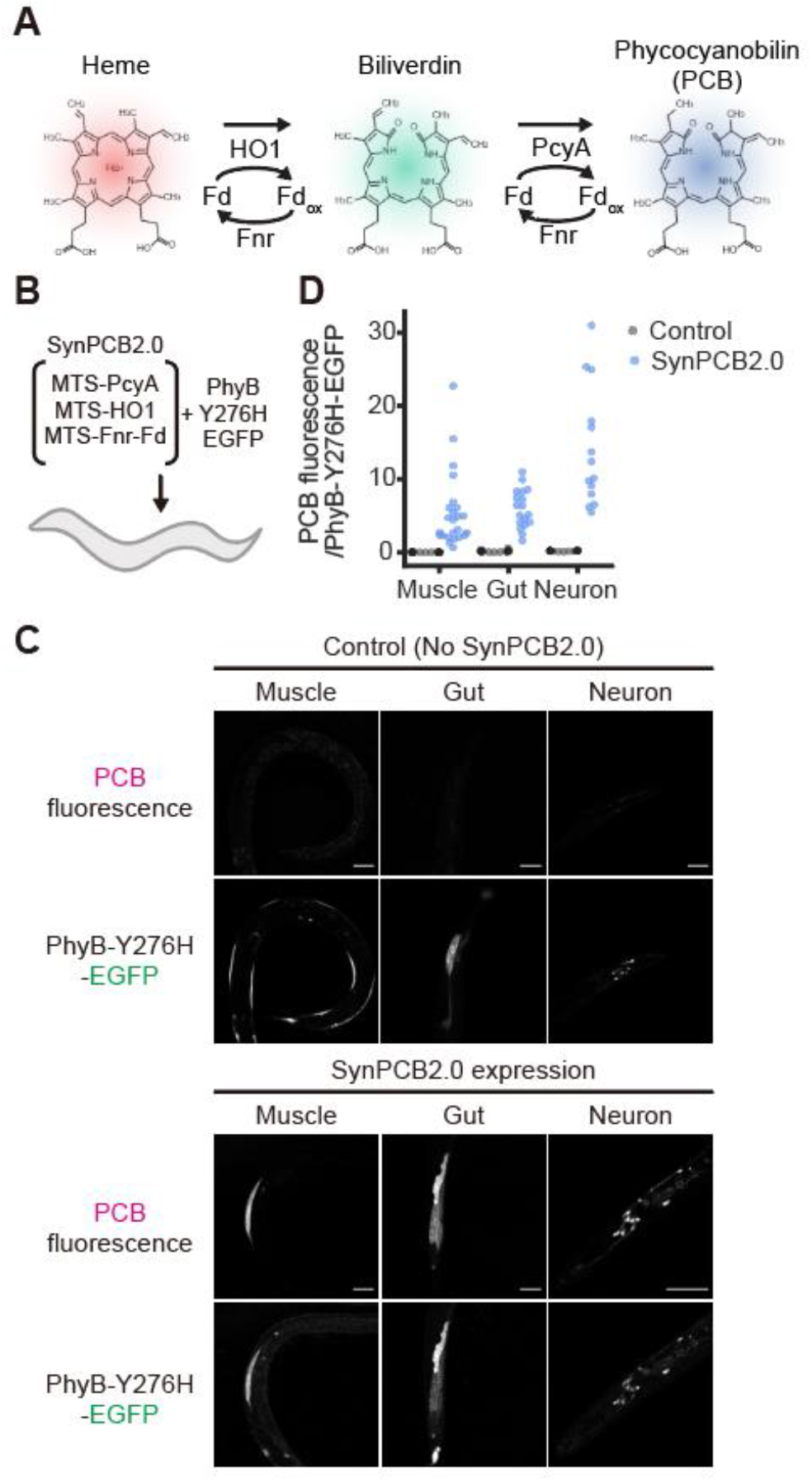
Efficient synthesis of PCB in various tissues of *C. elegans* by SynPCB2.0 expression. (A) Metabolisms of PCB synthesis. (B) Schematic representation of *C. elegans* expressing SynPCB2.0 and PhyB-Y276H-EGFP. (C) Representative images of PCB fluorescence (upper row) and PhyB-Y276H-EGFP (lower row) in control (upper panels) and SynPCB2.0-expressing worms (lower panels). Scale bar, 60 μm. (D) The ratio values of PCB fluorescence to PhyB-Y276H-EGFP fluorescence are plotted under the indicated conditions. Control: N = 19 (muscles), 19 (gut), and 21 (neurons). PCB addition: N = 24 (muscles), 18 (gut), 14 (neurons).

We examined PCB production by introducing SynPCB2.0 into *C. elegans*. To obtain the strains that express PhyB-Y276H-EGFP and SynPCB2.0 in multiple tissues, the worms expressing PhyB-Y276H-EGFP in the gut or neurons were interbred with the worms expressing SynPCB2.0 driven by a ubiquitous promoter, the *eft-3* promoter. The worms expressing PhyB-Y276H-EGFP in muscle were further injected with the plasmids encoding SynPCB2.0 induced by a muscle-specific promoter, the *myo-3* promoter. As expected, near-infrared fluorescence was hardly observed in worms without SynPCB2.0, while, in worms harboring SynPCB2.0, strong near-infrared fluorescence was observed in the muscle, gut, and neurons (Fig. 2C and 2D). These results indicate that the genetically encoded PCB synthesis system, SynPCB2.0, enables the synthesis of PCBs in various tissues of *C. elegans* such as the muscles, gut, and neurons.

### Reversible PhyB/PIF3 photoswitching with the SynPCB system in *C. elegans*

We next investigated whether the SynPCB system produces sufficient amounts of PCB for PhyB/PIF3 photoswitching. For this purpose, it is necessary to introduce at least three expression vectors, namely, PhyB, PIF3, and SynPCB, into *C. elegans*. To meet the requirement of simultaneously co-expressing different genes in the same tissues and to decrease potential toxicity by the ectopic expression of proteins in transgenic *C. elegans*, we introduced a Tet/Q hybrid system for the robust and drug-inducible control of transgene expression ^23^. The Tet/Q hybrid system comprises three components: a tetracycline responsive element (TRE) bearing the minimal promoter *Δpes-10*, a reverse TetR (rTetR) bearing the QF activation domain (QFAD), and a TetR fused to the PIE-1 repressor domain (Fig. 3A). In the absence of doxycycline (Dox), an analog of tetracycline, TetR-PIE blocks the transcription by inhibiting PolII carboxyl-terminal domain phosphorylation ^23,24^. Dox treatment induces gene expression through the transcriptional activation of rTetR-QFAD and the release of transcriptional inhibition of TetR-PIE.

**Figure 3.**
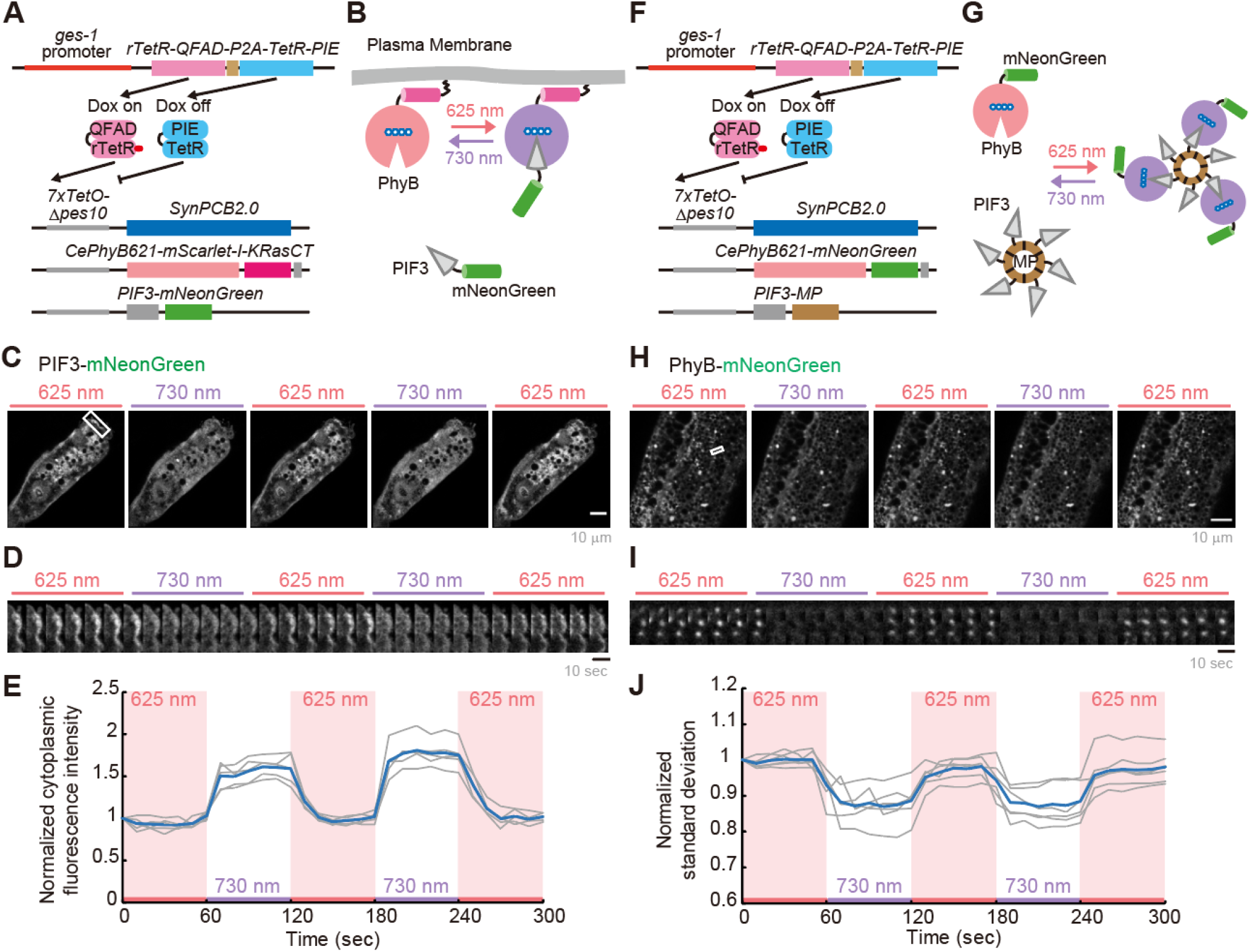
Light-induced photoswitching of PhyB/PIF3 with the SynPCB system in *C. elegans*. (A and F) Doxycycline (Dox)-inducible transgene expression system for the membrane translocation system (A) and aggregation formation system (F). (B and G) Schematic representation of the membrane translocation (B) and aggregation formation (G) through light-induced PhyB/PIF3 photoswitching. (C and D) The animals harboring the genes in panel A were treated with 1.0 ng/mL Dox for 3 days. The images were obtained every 10 sec by a confocal microscope with repeated red or far-red light exposure. Representative images of PIF3-mNeonGreen at the indicated time points (sec) are shown. The montage images in panel D highlight the membrane translocation of PIF3-mNeonGreen at the inset in panel C. Scale bar, 10 μm. (E) Fluorescence intensities of PIF3-mNeonGreen at the cytoplasm were normalized by the value at t = 0 and plotted as a function of elapsed time. Gray and blue lines represent the data extracted from one animal at the five different positions in two independent experiments and the averaged data, respectively. (H and I) The animals harboring the genes in panel F were treated with 1.0 ng/mL Dox for 2 days. The images were obtained every 10 sec by a confocal microscope with repeated red or far-red light exposure. Representative images of PhyB-mNeonGreen at the indicated time points (sec) are shown. The montage images in panel I highlight the PhyB-mNeonGreen aggregation at the inset in panel H. Scale bar, 10 μm. (J) Standard deviation (SD) values of the fluorescence intensities of PhyB-mNeonGreen at the cytoplasm were calculated and normalized by the value at t = 0. The normalized SD values are plotted as a function of elapsed time. Gray and blue lines represent the data extracted from four animals at five different positions in two independent experiments and averaged data, respectively.

To confirm the PhyB/PIF3 photoswitching, we implemented a system in which the binding of PIF3 and PhyB alters their subcellular localization. First, we localized PhyB at the plasma membrane and attempted to recruit PIF3 to the plasma membrane in a red light-dependent manner. In this system, rTetR-QFAD and TetR-PIE are expressed in the intestinal cells by the *ges-1* promoter (Fig. 3A). Since the intestinal cells are larger than other cells, we considered that intestinal cells would be suitable for observing changes in the subcellular localization of fluorescent proteins. In addition, the three plasmids were simultaneously introduced for expressing SynPCB2.0; PhyB 1-621 a.a. optimized for the codon usage of the *C. elegans* genome (CePhyB621) fused with mScarlet-I and a lipid modification site of the C-terminus of human K-Ras (CePhyB621-mScarlet-I-KRasCT); and PIF3-mNeonGreen. These proteins are expressed under the 7xTetO-*Δpes-10* promoter in the presence of 1.0 ng/mL Dox. Of note, a higher dose of Dox showed toxicity to worms (see Discussion). Red light illumination recruits PIF3-mNeonGreen from the cytoplasm to the plasma membrane through binding to CePhyB-mScarlet-I-KRasCT, and near-infrared light illumination induces dissociation of the PhyB-PIF3 complex, causing PIF3-mNeonGreen to return the cytoplasm (Fig. 3B). The animals harboring these genes were treated with 1.0 ng/mL Dox for 3 days, and the intestinal cells were observed. As we expected, PIF3-mNeonGreen rapidly and repeatedly changed its subcellular localization in a light-dependent manner (Fig. 3C-3E).

Next, we tested the photoswitching of PhyB/PIF3 using light-dependent molecular aggregation/dissolution. PIF3 fused with multimeric protein (MP), CePhyB621-mNeonGreen, and SynPCB2.0 are expressed in the intestinal cells in a Dox-inducible manner (Fig. 3F), showing aggregate formation when CePhyB621-mNeonGreen binds to PIF3-MP by red light illumination (Fig. 3G). In the transgenic animals treated with 10 ng/mL Dox for 2 days, CePhyB621-mNeonGreen rapidly and repeatedly formed aggregates in a light-dependent manner (Fig. 3H-3I). We evaluated the aggregate formation by the standard deviation of CePhyB621-mNeonGreen fluorescence intensity in intestinal cells and found that CePhyB621-mNeonGreen formed aggregates in a light-dependent manner multiple times (Fig. 3J). These results confirm our findings that the SynPCB2.0 system can produce sufficient amounts of PCB for photoswitching of PhyB/PIF3 in *C. elegans*.

### Light-induced manipulation of intracellular signaling with the PhyB/PIF3 and SynPCB system in *C. elegans*

Next, we attempted to manipulate intracellular signaling using the SynPCB system and PhyB/PIF3 photoswitching in *C. elegans*. It is well-known that cell signaling is triggered by the subcellular localization changes in signaling proteins, and therefore a chemical- or light-induced dimerization system allows manipulation of signaling pathways ^7,8,25^. Here, we introduced a system of Ca^2+^ release by red light, which has been demonstrated in cultured cells ^26^, into *C. elegans*. In this system, red light-induced binding of PIF3-Gαq/11 to membrane-anchored CePhyB621-mScarlet-I-KRasCT activates phospholipase Cβ (PLCβ) on the plasma membrane, resulting in an increase in cytosolic Ca^2+^ (Fig. 4A). PIF3-Gαq contains the constitutively active mutation, namely Q209L. We established the transgenic animals that express the SynPCB2.0, CePhyB621-mScarlet-I-KRasCT, PIF3-Gαq, and Ca^2+^ reporter GCaMP6s ^27^ in the gut in a drug-induced manner. The animals were treated with 100 ng/mL Dox for 3 days to express these proteins in the gut and illuminated with red or far-red light. We found a rapid increase in Ca^2+^ levels upon irradiation with red light (Fig. 4B). The Ca^2+^ increase induced by red light irradiation was repeatable and was rapidly suppressed by far-red light irradiation (Fig. 4B).

**Figure 4.**
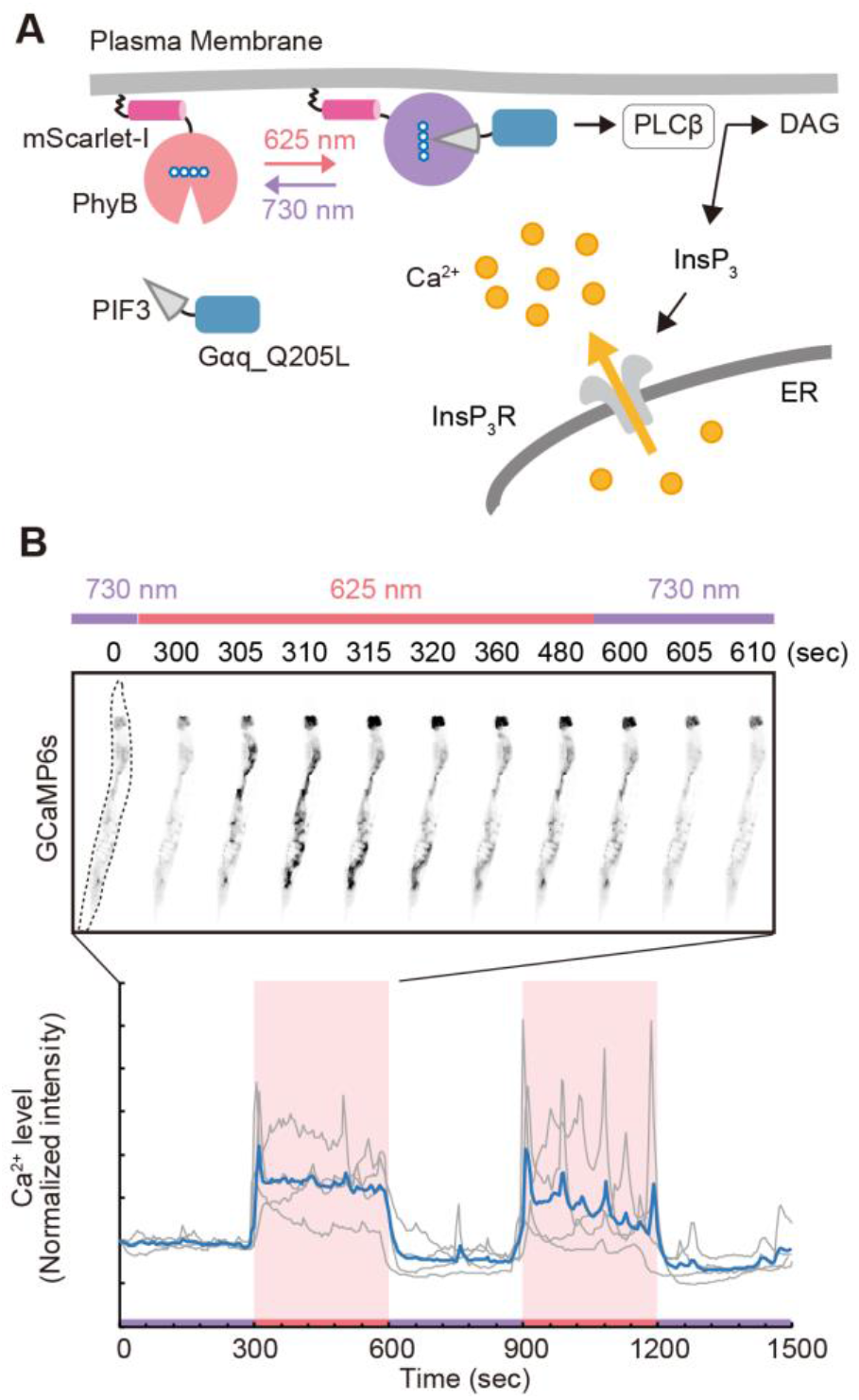
Light-induced increase in Ca2+ levels in *C. elegans* with the PhyB/PIF3 and SynPCB system. (A) Schematic illustration of light-induced manipulation of the Ca^2+^ level with the PhyB/PIF system. Upon red-light exposure, PIF3-Gαq is recruited to the plasma membrane through the binding to PhyB, leading to the production of diacylglycerol (DAG) and inositol 1,4,5-trisphosphate (InsP3). InsP3 binds to its receptor on the endoplasmic reticulum (ER) to induce Ca^2+^ release from ER. (B) The worms were treated with 100 ng/mL Dox for 3 days to express SynPCB2.0, CePhyB621-mScarlet-I-KRasCT, PIF3-Gα_q_, and GCaMP6s. They were fixed on beads and the Ca^2+^ level was observed upon the irradiation of red or far-red light. (Upper) Representative montage images of GCaMP6s in a single worm are shown with inverted image colors, *i*.*e*., black means high fluorescence intensity. (Lower) The normalized GCaMP6s intensity are plotted as a function of time. The gray and blue lines show individual and average results, respectively. N = 4 animals from the independent two experiments.

Similarly, we introduced a system in which ERK MAP kinase (Mpk-1) is activated by red light and ERK activation is visualized by a kinase translocation reporter (KTR) in *C. elegans* (Fig. S1A)^28,29^. The ERK-nKTR, which is optimized for *C. elegans*, revealed that ERK is activated by red light and inactivated by far-red light in the gut (Fig. S1B and 1C). Of note, some transgenic lines have high ERK activity and some do not respond to light (see Discussion). These results demonstrate that the SynPCB system and PhyB/PIF3 photoswitching enable the manipulation of intracellular signaling in *C. elegans* by red and far-red light.

### Induction of the defecation motor program in *C. elegans* by light

Finally, we tried to control the animals’ behavior with light, and focused on the defecation motor program (DMP) ^30^. DMP is a periodic behavior triggered approximately every 50 seconds during feeding (Fig. 5A). The first physical feature is the posterior body muscle contraction (pBoc), which is associated with the zigzag pattern of the intestinal lumen. Second, the worms show anterior body muscle contraction (aBoc), followed by enteric muscle contraction with expulsion (exp) of gut contents, and return to the intercycle ^31^. It has been shown that Ca^2+^ elevation in intestinal cells is required for DMP through PLCβ (egl-8) ^32^. Indeed, intestinal Ca2+ wave dynamics during DMP has been observed in *C. elegans* ^*33,34*^. On the other hand, it is unclear whether Ca^2+^ elevation of intestinal cells is sufficient to trigger DMP. For this purpose, we applied the red/far-red light-induced Ca^2+^ manipulation system shown in Figure 4 to DMP in *C. elegans*. To observe the morphology of a worm, a disposable 10 μL tip was cut out, a part of the tip was put on a thin agar plate, and an animal was placed inside the ring so that it would not escape from the field of view (Figs. 5B and 5C). At this time, since there was less food, the worm was starved, and thus autonomous DMP was suppressed. As expected, by switching from far-red light to red light, we often observed a zigzag pattern of the intestinal lumen, a typical pBoc phenotype, after a few seconds of red light illumination (Fig. 5D). The pBoc was triggered for a few seconds after Ca^2+^ elevation (Fig. 5E). We did not see any correlation between red light illumination and pBoc behavior in control nematodes lacking SynPCB2.0 (Fig. S2), negating the possibility that red light itself stimulates pBoc. These results indicate that the initial step of DMP, pBoc, is sufficiently triggered only by elevated Ca^2+^ in intestinal cells.

**Figure 5.**
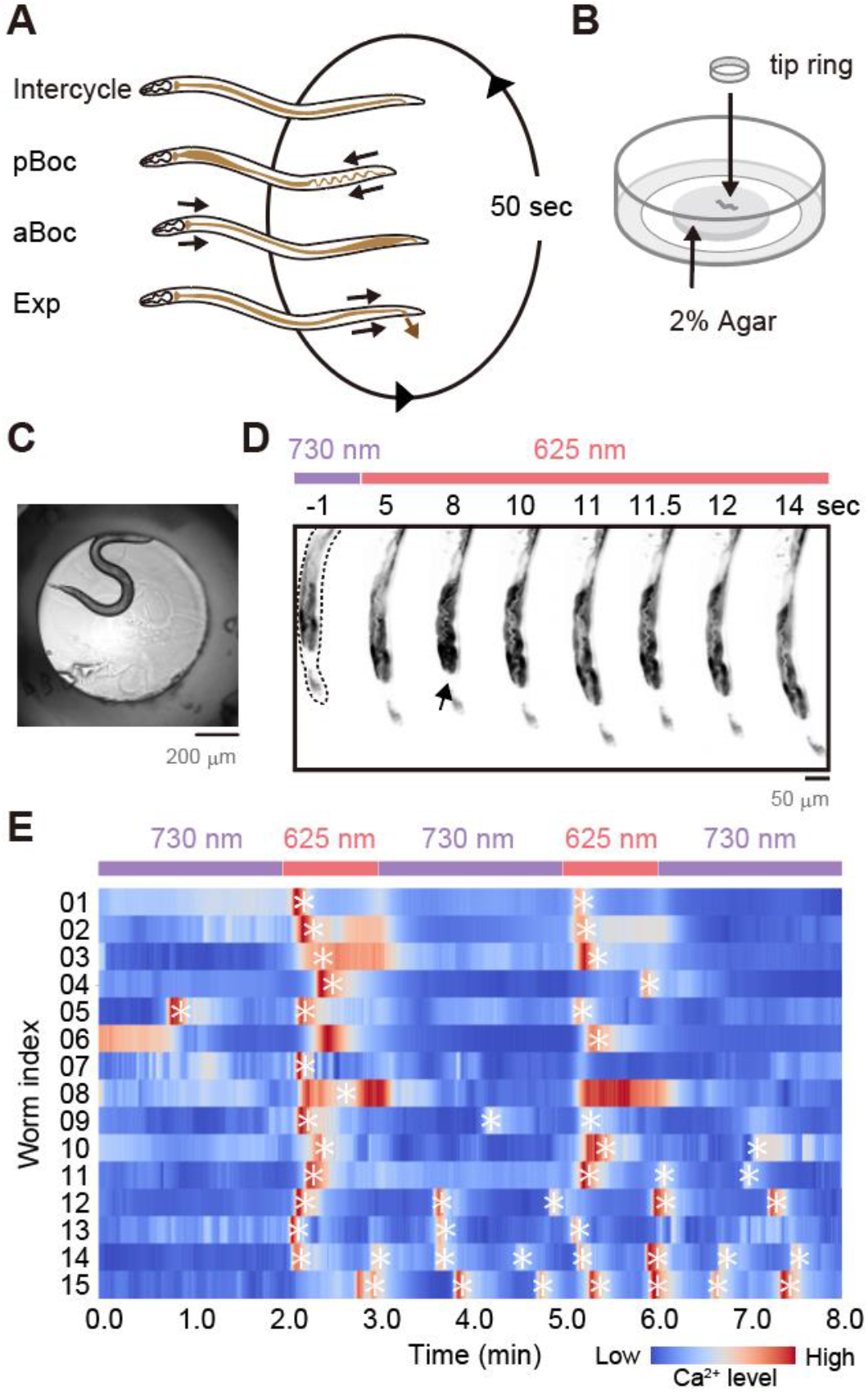
Light-induced Ca^2+^ elevation sufficiently induces an initial step of the defecation motor program (DMP) in *C. elegans*. (A) A schematic representation of the defecation motor program (DMP) in *C. elegans*: intercycle, pBoc (posterior body wall muscle contraction), aBoc (anterior body wall muscle contraction), and Exp (enteric muscle contraction with expulsion of gut contents). (B) An experimental setup for the observation of DMP. A tip ring made by cutting a disposable 10 μL tip was put on a thin agar plate, and a nematode was placed inside the ring. (C) A representative right field image. (D and E) The worms were treated with 100 ng/mL Dox for 3 days to express SynPCB2.0, CePhyB621-mScarlet-I-KRasCT, PIF3-Gαq, and GCaMP6s. They were placed in the tip ring shown in panel C, and the Ca^2+^ level was observed upon the irradiation of red or far-red light. (D) Representative montage images of GCaMP6s in a single worm are shown with inverted image colors. (E) The normalized GCaMP6s intensities in multiple nematodes (N = 15 worms) are shown as a heatmap. The asterisk indicates pBoc behavior.

## Discussion

In this study, we succeeded in introducing a genetically encoded PCB synthesis system, SynPCB, into *C. elegans*. One reason for the success in this model is the lack of a biliverdin reductase gene(s) in the *C. elegans* genome; biliverdin reductase catalyzes biliverdin and PCB to produce bilirubin and phycocyanorubin, respectively ^35^. The key to utilizing the PhyB-PIF system with the SynPCB system in mammalian cultured cells is the knockout or knockdown of the biliverdin reductase A (*blvra*) gene ^16^. Therefore, the absence of the biliverdin reductase gene in the *C. elegans* genome constitutes an advantage for the application of the SynPCB system.

One advantage of red light/far-red light optogenetics with the SynPCB system is that PCB can be ectopically synthesized in deeper tissues, where it is experimentally difficult to deliver exogenous PCB. Therefore, the SynPCB system obviates the need to purify and add PCB to living organisms. We showed that PCB does not reach the body muscle or nerves of *C. elegans* (Fig. 1). This is in contrast to the all-*trans* retinal, the chromophore of channelrhodopsin ^5^. Presumably because PCB is more than twice the size of all-*trans* retinal, PCB is less permeable to tissues than all-*trans* retinal. One advantage of adding purified chromophores is that it is not necessary to maintain the living organisms under the dark condition when not doing experiments. However, the introduction of the Dox-inducible system enables control of PCB synthesis and PhyB-PIF expression at the desired timing and expression levels, further facilitating the use of phytochrome-based optogenetics.

We directly demonstrated that Ca^2+^ elevation in the intestinal cells triggers pBoc, an initial step of DMP, based on the morphology of the gut lumen (Fig. 5). On the other hand, our observations did not conclusively determine whether the subsequent aBoc and exp steps were triggered. There are two possibilities: one is that we simply did not have enough knowledge to determine the aBoc and exp steps, and the other is that Ca^2+^ elevation in the intestinal cells is not, in itself, capable of inducing the steps after pBoc. Regarding the latter possibility, it has been reported that the increase in Ca^2+^ propagates like a wave from the posterior to anterior cells of the intestine ^33,34^. Because we globally illuminated the red/far-red light to the nematodes in the present study, intracellular Ca^2+^ levels rose almost simultaneously in cells throughout the intestine (Fig. 4). For this reason, it is likely that the later steps of DMP, such as aBoc and exp, require temporally and spatially coordinated Ca^2+^ waves, as observed in actual *C. elegans*. In addition to this, considering that we illuminated red/far-red light under starvation conditions to inhibit spontaneous DMP, other stimuli from the intestinal lumen may be required for the induction of aBoc and exp.

Two drawbacks of implementing the SynPCB system should be mentioned. The first is that SynPCB expression induces toxicity. We established multiple transgenic strains that constantly express the genes for PCB synthesis (Fig. 2), but the fraction of worms showing PCB synthesis was generally not high. We also introduced a drug-inducible expression system and investigated the optimal concentration of Dox and found that high doses of Dox showed toxicity. These findings suggest that SynPCB is toxic to *C. elegans* when expressed at high levels. Previously, we had applied SynPCB to cells widely used in cultures, such as HeLa cells, mouse embryonic stem cells, fission yeast, and *Xenopus* embryos ^17,19^, but *C. elegans* are the first for which SynPCB is toxic. Since *C. elegans* is unable to synthesize Heme on its own and takes it in from its diet ^36^, the expression of the SynPCB system might be associated with toxicity. In any case, care should be taken in applying the SynPCB system in other living organisms. The second issue is the control of stoichiometry. In this experimental system, we introduced multiple genes into *C. elegans*: examples include SynPCB, photoswitchable proteins (e.g., PhyB-PIF system), a reporter (e.g., GCaMP), and a drug-inducible unit (Figs. 4, 5, and S1). The stoichiometry between PhyB and PIF is crucial for the optical manipulation of the PhyB-PIF system ^17^. Although, as mentioned above, excessive expression of the SynPCB system causes toxicity, low PCB synthesis is incapable of optically manipulating the PhyB-PIF switch in the first place. Therefore, it is necessary to optimize the expression levels of each gene.

Finally, we would like to mention other potential applications of SynPCB; the SynPCB system could be used not only for phytochrome-based optogenetic tools but also for near-infrared fluorescent proteins that bind to linear tetrapyrroles. Recently, we have shown that the application of the SynPCB system in fission yeast allows for multicolor fluorescence imaging using the near-infrared fluorescent protein iRFP ^37^. Combined with knockout of the Blvra gene ^38^, the SynPCB system would provide opportunities for bright near-infrared fluorescence imaging in large animals.

## Materials and methods

### Plasmids

All plasmids used in this study are listed in Supplementary Information Table S1 with Benchling links, which include the sequences and plasmid maps. The cDNAs were subcloned into vectors through conventional ligation with Ligation high Ver.2 (Toyobo, Japan) or NEBuilder HiFi DNA Assembly (New England Biolabs, MA) according to the manufacturer’s instructions. The cDNAs of SynPCB2.0 (*PcyA, HO1*, and *Fd-Fnr* chimera genes of *Thermosynechococcus elongatus* BP-1), Gαq (*egl-30*)_SS_Q205L and GCaMP6s were synthesized with codon-optimization for *C. elegans* genome by FASMAC (Kanagawa, Japan) ^17,19^. *lin-44p::gfp* was a kind gift from Dr. De Bono (ISTA, Austria). TRE-Δpes-10 and rtetR-QFAD-P2A-tetR-pie1 were amplified from TC358 TRE-*Δpes-10*::GFP and TC374 *rpl-28p*::rtTA(Q)(rtetR-QFAD)::P2A::mKate::T2A::tTS (tetR-pie-1), both of which were a kind gift of Dr. Chi (ShanghaiTech University, China) ^23^. *ges-1* promoter was a kind gift from Dr. Iino (The University of Tokyo, Japan).

### Strains and transformation

All strains established in this study are listed in the Supplementary Information Table S2. Animals were grown at 15°C or 23°C under standard conditions on Nematode Growth Medium (NGM) seeded with *Escherichia coli* OP50. All transgenic lines carrying extrachromosome arrays were generated on the N2 background using standard protocols ^39^. To externally add PCB, 10 uL of purified 5 mM PCB (SC-1800; SciChem, Germany) dissolved in DMSO was dropped onto the lawn of *E. coli* OP50 and then the animals were fed overnight with food containing PCB or only DMSO. Doxycycline (D45115; Tokyo Chemical Industry, Japan) was dissolved in DMSO (10 mg/mL), and the resulting solution was diluted with M9 buffer to get 1 ng/uL stock solution. Before use, the stock was diluted to 1 ∼20 pg/uL with M9 buffer, and 200 uL was added onto each nematode growth medium (NGM) plate (IWAKI: 35 mm) seeded with OP50. L2/L3 larvae were transferred to plates and exposed to doxycycline for 2 to 4 days at 23°C.

### Fluorescence imaging

Worms were immobilized by 0.5 M NaN3 treatment (Figs. 1 and 2) or polystyrene nanoparticles (2.5% by volume, 0.1 μm diameter, Polysciences) (Figs. 3-4) ^40^. For the immobilization with polystyrene nanoparticles, doxycycline-exposed worms were placed in a suspension of polystyrene nanoparticles on an agarose pad containing 8-10% agarose in M9 buffer, and then the worms were covered with a coverslip. To observe moving worms, the doxycycline-exposed worms were placed on a 2% agar pad on a 35 mm glass-base dish (IWAKI, Japan). Then, a P10 micropipette-tip was cut into small rings, which were placed around the worms on the agar pad to limit their migration.

For confocal fluorescence imaging (Figs. 1-3), worms were imaged with a TCS SP5 microscope (Leica Microsystems, Germany) using an oil immersion objective lens (HCX PL APO 63/x1.4-0.6 oil; Leica Microsystems). The excitation laser and fluorescence filter settings were as follows: excitation laser, 488 nm (mNeonGreen) and 543 nm (mScarlet-I); dichroic mirror, TD 488/543/633 dichroic mirror; detector, HyD 520-590 nm (mNeonGreen) and HyD 600-650 nm (mScarlet-I). LEDs for illumination with red (625 nm) and far-red (735 nm) light were purchased from Optocode and controlled manually.

For Ca^2+^ imaging (Figs. 4 and 5), worms were imaged with an IX83 inverted microscope (Olympus, Japan) equipped with an sCMOS camera (ORCA-528 Fusion BT; Hamamatsu Photonics, Japan), an oil objective lens (UPLXAPO40X, NA 0.95; Olympus) or a dry objective lens (UPLXAPO10X, NA; Olympus), and a spinning disk confocal unit (CSU-W1; Yokogawa Electric Corporation, Japan). The excitation laser and fluorescence filter settings were as follows: excitation laser, 488 nm for GCaMP6s; dichroic mirror, DM405/488/561/640; emission filters, 500-550 nm for GCaMP6s (Yokogawa Electric).

### Imaging analysis

All fluorescence imaging data were analyzed and quantified with Fiji (Image J)^41^. For all images, the background was subtracted by the rolling-ball method and images were registered by StackReg, a Fiji plugin to correct misregistration, if needed. To quantify the changes in cytoplasmic fluorescence intensity (Fig. 3E), the ROIs were manually selected and normalized by the mean fluorescence intensity of the first image. To obtain the normalized standard deviation (Fig. 3J), the standard deviation of fluorescence intensity in whole worms was quantified and normalized by the standard deviation value of the first image. To quantify the Ca^2+^ levels (Figs. 4 and 5), the mean fluorescence intensity of GCaMP6s in whole worms was measured and normalized by the mean fluorescence intensity of the first image. The timing of worm expulsion was identified by the naked eye.

## Acknowledgments

We thank all members of the Aoki Laboratory for their helpful discussions and assistance. S.O. was supported by The Sumitomo Foundation (Basic Science Research Projects). K.A. was supported by a CREST, JST Grant (JPMJCR1654) and JSPS KAKENHI Grants (nos. 18H04754 “Resonance Bio”, 19H05798, and 22H02625). K.D.K. was supported by JSPS KAKNHI Grants (nos. 21H02533, 21H05299) and Joint Research of the Exploratory Research Center on Life and Living Systems (ExCELLS) (ExCELLS program No. 22EXC206).

## Author contributions

S.O. and K.A. conceptualized the study. S.O., K.D.K., A.N., and K.A. designed the optogenetic tools. S.O. and E.E. performed all experiments and analyzed the data. S.O., E.E., A.N., K.D.K., and K.A. wrote the manuscript.

## References

(1) Wood, W. B. The Nematode Caenorhabditis Elegans; Cold Spring Harbor Laboratory, 1988.

(2) Inoue, T.; Heo, W. D.; Grimley, J. S.; Wandless, T. J.; Meyer, T. An Inducible Translocation Strategy to Rapidly Activate and Inhibit Small GTPase Signaling Pathways. Nat. Methods 2005, 2 (6), 415–418.

(3) Inoue, T.; Meyer, T. Synthetic Activation of Endogenous PI3K and Rac Identifies an AND-Gate Switch for Cell Polarization and Migration. PLoS One 2008, 3 (8), e3068.

(4) Fenno, L.; Yizhar, O.; Deisseroth, K. The Development and Application of Optogenetics. Annu. Rev. Neurosci. 2011, 34 (1), 389–412.

(5) Nagel, G.; Brauner, M.; Liewald, J. F.; Adeishvili, N.; Bamberg, E.; Gottschalk, A. Light Activation of Channelrhodopsin-2 in Excitable Cells of Caenorhabditis Elegans Triggers Rapid Behavioral Responses. Curr. Biol. 2005, 15 (24), 2279–2284.

(6) Bergs, A.; Schultheis, C.; Fischer, E.; Tsunoda, S. P.; Erbguth, K.; Husson, S. J.; Govorunova, E.; Spudich, J. L.; Nagel, G.; Gottschalk, A.; Liewald, J. F. Rhodopsin Optogenetic Toolbox v2.0 for Light-Sensitive Excitation and Inhibition in Caenorhabditis Elegans. PLoS One 2018, 13 (2), e0191802.

(7) Goto, Y.; Kondo, Y.; Aoki, K. Visualization and Manipulation of Intracellular Signaling. Adv. Exp. Med. Biol. 2021, 1293, 225–234.

(8) Farahani, P. E.; Reed, E. H.; Underhill, E. J.; Aoki, K.; Toettcher, J. E. Signaling, Deconstructed: Using Optogenetics to Dissect and Direct Information Flow in Biological Systems. Annu. Rev. Biomed. Eng. 2021. https://doi.org/10.1146/annurev-bioeng-083120-111648.

(9) Endo, M.; Hattori, M.; Toriyabe, H.; Ohno, H.; Kamiguchi, H.; Iino, Y.; Ozawa, T. Optogenetic Activation of Axon Guidance Receptors Controls Direction of Neurite Outgrowth. Sci. Rep. 2016, 6, 23976.

(10) Fielmich, L.-E.; Schmidt, R.; Dickinson, D. J.; Goldstein, B.; Akhmanova, A.; van den Heuvel, S. Optogenetic Dissection of Mitotic Spindle Positioning in Vivo. Elife 2018, 7. https://doi.org/10.7554/eLife.38198.

(11) De Henau, S.; Pagès-Gallego, M.; Pannekoek, W.-J.; Dansen, T. B. Mitochondria-Derived H2O2 Promotes Symmetry Breaking of the C. Elegans Zygote. Dev. Cell 2020, 53 (3), 263–271.e6.

(12) Deng, W.; Bates, J. A.; Wei, H.; Bartoschek, M. D.; Conradt, B.; Leonhardt, H. Tunable Light and Drug Induced Depletion of Target Proteins. Nat. Commun. 2020, 11 (1), 304.

(13) Gong, J.; Yuan, Y.; Ward, A.; Kang, L.; Zhang, B.; Wu, Z.; Peng, J.; Feng, Z.; Liu, J.; Xu, X. Z. S. The C. Elegans Taste Receptor Homolog LITE-1 Is a Photoreceptor. Cell 2016, 167 (5), 1252– 1263.e10.

(14) Shimizu-Sato, S.; Huq, E.; Tepperman, J. M.; Quail, P. H. A Light-Switchable Gene Promoter System. Nat. Biotechnol. 2002, 20 (10), 1041–1044.

(15) Levskaya, A.; Weiner, O. D.; Lim, W. A.; Voigt, C. A. Spatiotemporal Control of Cell Signalling Using a Light-Switchable Protein Interaction. Nature 2009, 461 (7266), 997–1001.

(16) Müller, K.; Engesser, R.; Timmer, J.; Nagy, F.; Zurbriggen, M. D.; Weber, W. Synthesis of Phycocyanobilin in Mammalian Cells. Chem. Commun. 2013, 49 (79), 8970.

(17) Uda, Y.; Goto, Y.; Oda, S.; Kohchi, T.; Matsuda, M.; Aoki, K. Efficient Synthesis of Phycocyanobilin in Mammalian Cells for Optogenetic Control of Cell Signaling. Proceedings of the National Academy of Sciences 2017, 114 (45), 11962–11967.

(18) Kyriakakis, P.; Catanho, M.; Hoffner, N.; Thavarajah, W.; Hu, V. J.; Chao, S.-S.; Hsu, A.; Pham, V.; Naghavian, L.; Dozier, L. E.; Patrick, G. N.; Coleman, T. P. Biosynthesis of Orthogonal Molecules Using Ferredoxin and Ferredoxin-NADP+ Reductase Systems Enables Genetically Encoded PhyB Optogenetics. ACS Synth. Biol. 2018, 7 (2), 706–717.

(19) Uda, Y.; Miura, H.; Goto, Y.; Yamamoto, K.; Mii, Y.; Kondo, Y.; Takada, S.; Aoki, K. Improvement of Phycocyanobilin Synthesis for Genetically Encoded Phytochrome-Based Optogenetics. ACS Chem. Biol. 2020, 15 (11), 2896–2906.

(20) Yang, X.; Jost, A. P.-T.; Weiner, O. D.; Tang, C. A Light-Inducible Organelle-Targeting System for Dynamically Activating and Inactivating Signaling in Budding Yeast. Mol. Biol. Cell 2013, 24 (15), 2419–2430.

(21) Beyer, H. M.; Juillot, S.; Herbst, K.; Samodelov, S. L.; Müller, K.; Schamel, W. W.; Römer, W.; Schäfer, E.; Nagy, F.; Strähle, U.; Weber, W.; Zurbriggen, M. D. Red Light-Regulated Reversible Nuclear Localization of Proteins in Mammalian Cells and Zebrafish. ACS Synth. Biol. 2015, 4 (9), 951–958.

(22) Buckley, C. E.; Moore, R. E.; Reade, A.; Goldberg, A. R.; Weiner, O. D.; Clarke, J. D. W. Reversible Optogenetic Control of Subcellular Protein Localization in a Live Vertebrate Embryo. Dev. Cell 2016, 36 (1), 117–126.

(23) Mao, S.; Qi, Y.; Zhu, H.; Huang, X.; Zou, Y.; Chi, T. A Tet/Q Hybrid System for Robust and Versatile Control of Transgene Expression in C. Elegans. iScience 2019, 11, 224–237.

(24) Ghosh, D.; Seydoux, G. Inhibition of Transcription by the Caenorhabditis Elegans Germline Protein PIE-1: Genetic Evidence for Distinct Mechanisms Targeting Initiation and Elongation. Genetics 2008, 178 (1), 235–243.

(25) DeRose, R.; Miyamoto, T.; Inoue, T. Manipulating Signaling at Will: Chemically-Inducible Dimerization (CID) Techniques Resolve Problems in Cell Biology. Pflugers Arch. 2013, 465 (3), 409–417.

(26) Yu, G.; Onodera, H.; Aono, Y.; Kawano, F.; Ueda, Y.; Furuya, A.; Suzuki, H.; Sato, M. Optical Manipulation of the Alpha Subunits of Heterotrimeric G Proteins Using Photoswitchable Dimerization Systems. Sci. Rep. 2016, 6 (1), 35777.

(27) Chen, T.-W.; Wardill, T. J.; Sun, Y.; Pulver, S. R.; Renninger, S. L.; Baohan, A.; Schreiter, E. R.; Kerr, R. A.; Orger, M. B.; Jayaraman, V.; Looger, L. L.; Svoboda, K.; Kim, D. S. Ultrasensitive Fluorescent Proteins for Imaging Neuronal Activity. Nature 2013, 499 (7458), 295–300.

(28) Aoki, K.; Kumagai, Y.; Sakurai, A.; Komatsu, N.; Fujita, Y.; Shionyu, C.; Matsuda, M. Stochastic ERK Activation Induced by Noise and Cell-to-Cell Propagation Regulates Cell Density-Dependent Proliferation. Mol. Cell 2013, 52 (4), 529–540.

(29) de la Cova, C.; Townley, R.; Regot, S.; Greenwald, I. A Real-Time Biosensor for ERK Activity Reveals Signaling Dynamics during C. Elegans Cell Fate Specification. Dev. Cell 2017, 42 (5), 542–553.e4.

(30) Liu, D. W.; Thomas, J. H. Regulation of a Periodic Motor Program in C. Elegans. J. Neurosci. 1994, 14 (4), 1953–1962.

(31) Branicky, R.; Hekimi, S. What Keeps C. Elegans Regular: The Genetics of Defecation. Trends Genet. 2006, 22 (10), 571–579.

(32) Lackner, M. R.; Nurrish, S. J.; Kaplan, J. M. Facilitation of Synaptic Transmission by EGL-30 Gqalpha and EGL-8 PLCbeta: DAG Binding to UNC-13 Is Required to Stimulate Acetylcholine Release. Neuron 1999, 24 (2), 335–346.

(33) Teramoto, T.; Iwasaki, K. Intestinal Calcium Waves Coordinate a Behavioral Motor Program in C. Elegans. Cell Calcium 2006, 40 (3), 319–327.

(34) Nehrke, K.; Denton, J.; Mowrey, W. Intestinal Ca2+ Wave Dynamics in Freely Moving C. Elegans Coordinate Execution of a Rhythmic Motor Program. Am. J. Physiol. Cell Physiol. 2008, 294 (1), C333–C344.

(35) Terry, M. J.; Maines, M. D.; Lagarias, J. C. Inactivation of Phytochrome- and Phycobiliprotein-Chromophore Precursors by Rat Liver Biliverdin Reductase. J. Biol. Chem. 1993, 268 (35), 26099–26106.

(36) Rao, A. U.; Carta, L. K.; Lesuisse, E.; Hamza, I. Lack of Heme Synthesis in a Free-Living Eukaryote. Proc. Natl. Acad. Sci. U. S. A. 2005, 102 (12), 4270–4275.

(37) Sakai, K.; Kondo, Y.; Fujioka, H.; Kamiya, M.; Aoki, K.; Goto, Y. Near-Infrared Imaging in Fission Yeast Using a Genetically Encoded Phycocyanobilin Biosynthesis System. J. Cell Sci. 2021, 134 (24). https://doi.org/10.1242/jcs.259315.

(38) Kobachi, K.; Kuno, S.; Sato, S.; Sumiyama, K.; Matsuda, M.; Terai, K. Biliverdin Reductase-A Deficiency Brighten and Sensitize Biliverdin-Binding Chromoproteins. Cell Struct. Funct. 2020, 45 (2), 131–141.

(39) Mello, C.; Fire, A. DNA Transformation. Methods Cell Biol. 1995, 48, 451–482.

(40) Kim, E.; Sun, L.; Gabel, C. V.; Fang-Yen, C. Long-Term Imaging of Caenorhabditis Elegans Using Nanoparticle-Mediated Immobilization. PLoS One 2013, 8 (1), e53419.

(41) Schindelin, J.; Arganda-Carreras, I.; Frise, E.; Kaynig, V.; Longair, M.; Pietzsch, T.; Preibisch, S.; Rueden, C.; Saalfeld, S.; Schmid, B.; Tinevez, J. Y.; White, D. J.; Hartenstein, V.; Eliceiri, K.; Tomancak, P.; Cardona, A. Fiji: An Open-Source Platform for Biological-Image Analysis. Nat. Methods 2012, 9 (7), 676–682.

