## Supplementary Information for "Optogenetic control of cell signaling with red/far-red light-responsive optogenetic tools in *Caenorhabditis elegans*"

**Supplementary Figure 1**

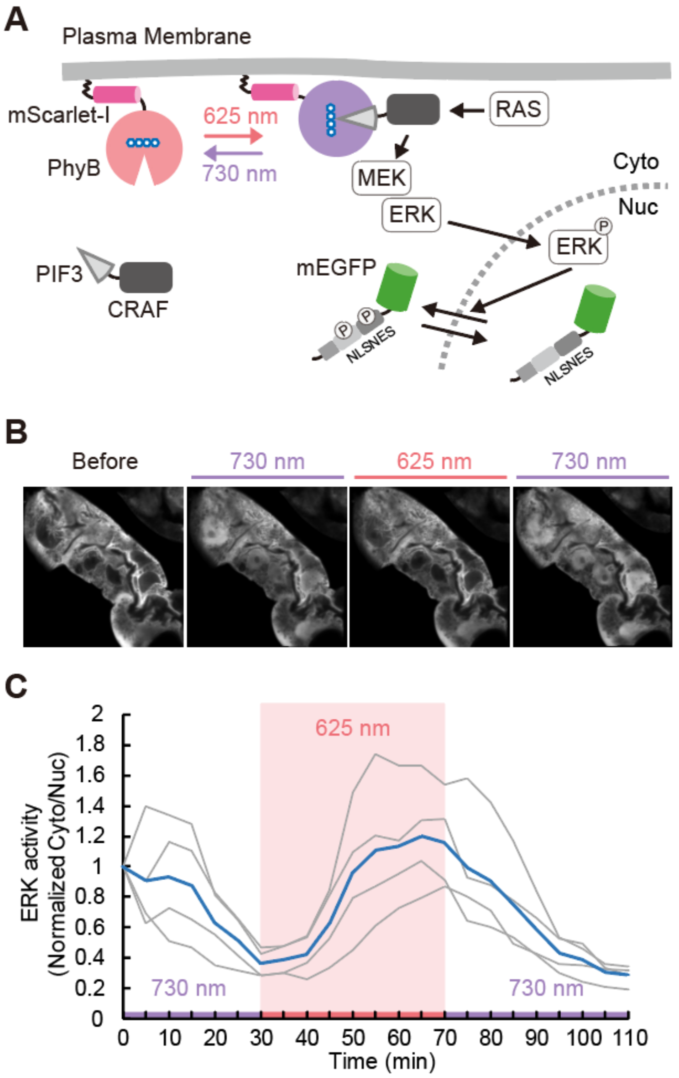


**Figure S1. Optogenetic ERK activation in gut cells of *C. elegans*.**

(A) Schematic illustration of light-induced ERK activation with the PhyB/PIF system. Upon red-light exposure, PIF3-CRAF is recruited to the plasma membrane through the binding to PhyB, leading to the activation of MEK and ERK. Upon phosphorylation by ERK, ERK-KTR-mEGFP (kinase translocation reporter) is translocated from nucleus to cytoplasm. (B) The worms were treated with 5 ng/mL Dox for 2 days to express SynPCB2.0, CePhyB621-mScarlet-I-HRasCT, PIF3-CRAF, and ERK-KTR-mEGFP. They were fixed on beads and ERK-KTR-mEGFP was observed upon the irradiation of red or far-red light. Representative montage images of ERK-KTR-mEGFP in a single worm are shown. (C) The cytoplasm-to-nucleus (Cyt/Nuc) ratio of ERK-KTR-mEGFP was quantified and normalized by the first time point. The normalized Cyt/Nuc ratio is plotted as a function of time. The gray and blue lines show individual and average results, respectively. N = 4 cells in panel B. We performed seven independent experiments and observed 19 worms. Among them, four worms showed light-induced localization change in ERK-KTR-mEGFP.

**Supplementary Figure 2**

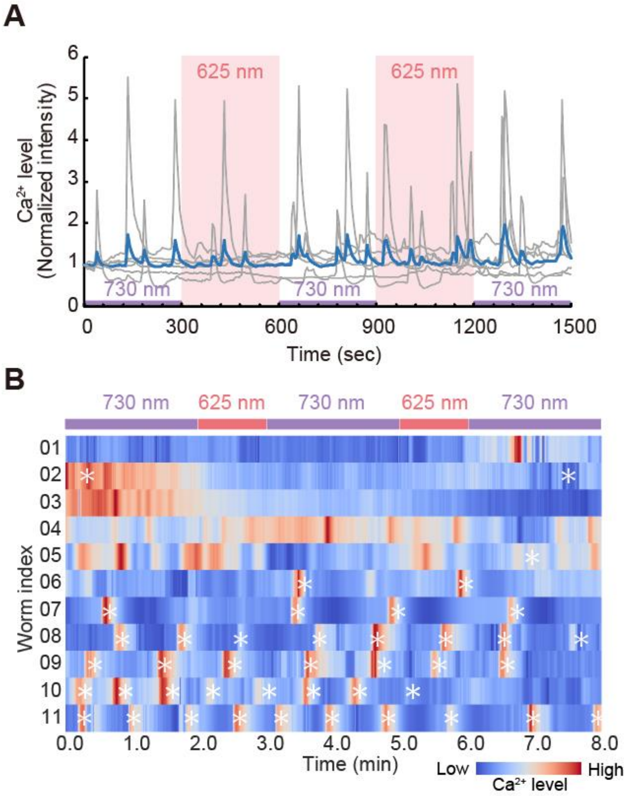


**Figure S2. Negative control experiments for $Ca^{2+}$ levels upon red and far-red light illumination.** (A) The worms were treated with 100 ng/mL Dox for 3 days to express CePhyB621-mScarlet-I-KRasCT, PIF3-Gαq, and GCaMP6s without SynPCB2.0. They were fixed on beads and $Ca^{2+}$ level was observed upon the irradiation of red or far-red light. The normalized GCaMP6s intensity are plotted as a function of time. The gray and blue lines show individual and average results, respectively. N = 6. (B) The worms were treated with 100 ng/mL Dox for 3 days to express CePhyB621-mScarlet-I-KRasCT, PIF3-$G\alpha_q$, and GCaMP6s without SynPCB2.0. They were placed in tip rings as shown in Figure 5C, and the $Ca^{2+}$ level was observed upon the irradiation of red or far-red light. The normalized GCaMP6s intensity in multiple worms (N = 11 worms from three independent experiments) is shown as a heatmap. The asterisk indicates pBoc behavior.
